# Exploring GNRA tetraloop-like motifs in nucleic acid 3D structures

**DOI:** 10.1101/2025.07.03.663028

**Authors:** Janusz M. Bujnicki, Eugene F. Baulin

## Abstract

Structured nucleic acids play key roles in a variety of molecular mechanisms and have numerous applications in medicine and biotechnology. Stem-loops with specific tetraloop sequences, such as UNCG and GNRA, are among the most well-studied recurrent building blocks of structured RNAs, known to adopt very stable structures. The GNRA tetraloop motif frequently engages in long-range interactions with specific RNA receptors and can also be recognized by proteins. While several examples of GNRA-like conformations have been reported in non-tetraloop strands, no systematic survey has been conducted to explore the landscape of these variants. In this work, we report a comprehensive survey of GNRA tetraloop-like motifs within known 3D structures of nucleic acids, without imposing restraints on sequence, loop type, backbone topology, or interactions involved. We identified twelve recurrent backbone topology variants of the motif, including all five possible two-strand variants and six three-strand variants. Among the pentaloop variants, we observed four different locations of the looped-out residues, with three of them directly interacting with proteins. These GNRA-like motifs highlight the versatility of favorable RNA 3D conformations to fit in diverse backbone contexts and modulate intermolecular interactions, which can be leveraged in RNA motif design.

## INTRODUCTION

Structured nucleic acids play key roles in a variety of molecular mechanisms [1-3] and have numerous applications in medicine and biotechnology [4, 5]. The 3D structure of nucleic acids is organized hierarchically, comprising recurrent building blocks such as base pairs, helical segments, and tertiary motifs, with tetraloops of specific sequence patterns among the most prevalent [6-8]. GNRA tetraloops (N stands for aNy nucleotide, R stands for puRine) are overrepresented in non-coding RNAs, including ribosomal RNAs, ribozymes, and riboswitches [8, 9]. They adopt very stable structures in both RNA and DNA variants [10].

The characteristic structural features of the GNRA tetraloop motif include the trans-Sugar-Hoogsteen (tSH) G-A base pair, G-RpA base-phosphate hydrogen bond, R-rG base-ribose hydrogen bond, and NpR-G oxygen-base stacking [11] (Figure 1). The U-turn motif (GNR) of the GNRA tetraloop arranges the stacked loop bases such that their Watson-Crick (WC) edges remain available for interactions [12, 13]. Consequently, GNRA tetraloops often form long-range tertiary interactions, primarily through A-minor motifs [14-17], for example, interactions between the L9 loop and the P5 helix of a group I intron required for its proper folding and self-splicing activity [15]. The highly specific GAAA tetraloop/11nt receptor interaction is among the most stable long-range tertiary motifs formed with a GNRA loop [17] and has been widely used as a building block in RNA nanostructures and scaffolding [16, 18]. Conserved GNRA tetraloops within viral internal ribosome entry site (IRES) elements are essential for their role in translation [19-21]. The universally conserved sarcin-ricin loop (SRL) of the ribosome contains a GAGA tetraloop that interacts with the elongation factors EF-Tu and EF-G during translation and is specifically recognized by the ricin toxin [22, 23].

**Figure 1.**
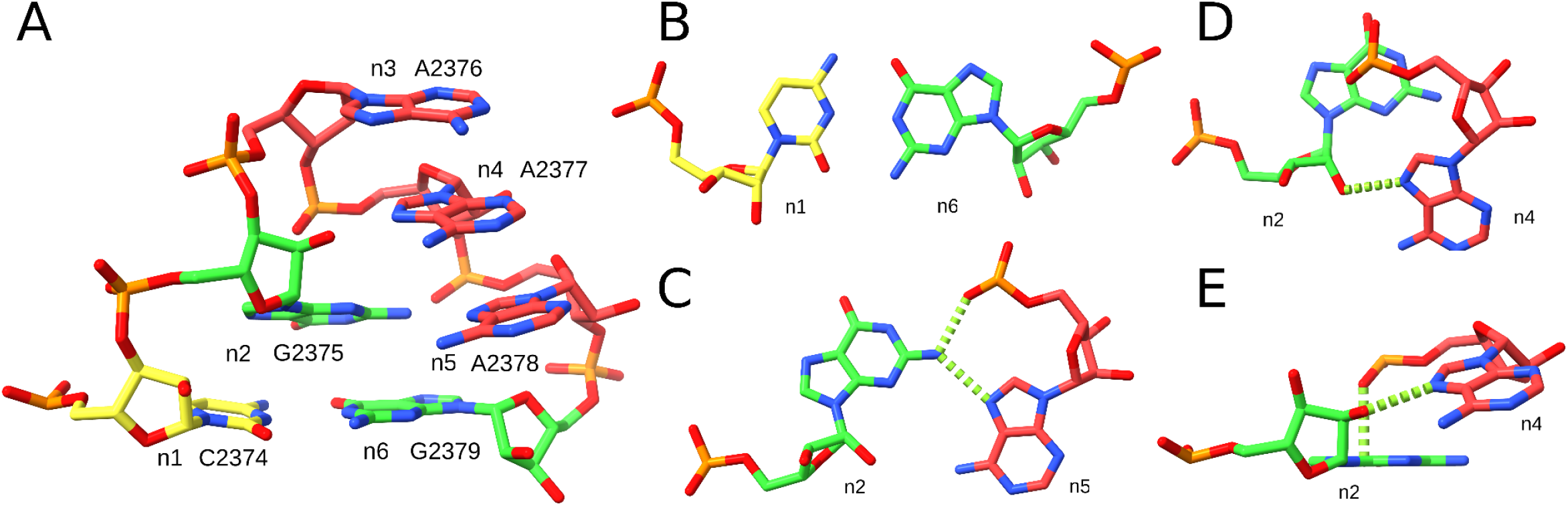
Reference instance of the GNRA tetraloop motif. (**A**) Residues n1-n6 (PDB entry 8VTW, 23S rRNA, chain 1A, residues 2374-2379), (**B**) Flanking canonical C-G base pair between residues n1 and n6. (**C**) tSH G-A base pair between residues n2 and n5, and the G-RpA base-phosphate interaction. (**D**) Top and (**E**) side views of the R-rG base-ribose hydrogen bond and the NpR-G oxygen-base stacking interaction between residues n2 and n4. The color scheme shows guanosines in green, adenosines in red, cytidines in yellow, and uridines in blue (not present in this figure). Dashed green lines indicate key interactions. The figure was prepared using UCSF ChimeraX (https://www.cgl.ucsf.edu/chimerax/).

GNRA-like motifs can also be formed by non-tetraloop strands, such as pentaloops [24-26] and hexaloops [27]. These non-tetraloop variants can also be specifically recognized by proteins, such as in the N-peptide-BoxB RNA complexes of bacteriophages [24-29]. It has been shown that the GNRA sequence can be split and interrupted by other nucleotides (GN(n)/RA) and still adopt the characteristic GNRA structure capable of forming long-range interactions [30, 31]. The UAA/GAN internal loop, found in 23S rRNA, RNase P, and group I and II introns [32], falls within this extended consensus (GAN/UAA in reverse order of strands) and is also considered a GNRA-tetraloop-like motif [33, 34]. Similar to GNRA-tetraloop/receptor motifs, UAA/GAN internal loops form cross-strand AAA stacks involved in long-range A-minor interactions [32]. GNRA-like motifs can even mediate RNA-ligand interactions, for example, in the complex of an in vitro selected ATP-binding aptamer with AMP, where the ligand intercalates beneath the R residue of the GNR strand [35]. Moreover, GNRA-like motifs can be formed by non-GNRA sequences [17, 36, 37], such as UMAC loops (where M is A or C), found in ribosomes and BoxB RNAs of bacteriophages [27, 36, 38]. While several previous studies have surveyed the structural landscape of tetraloops and larger hairpins [36, 39], the GNRA tetraloop motif has not been explored in a backbone topology-independent manner. As a result, the diversity of known GNRA-like motif topologies remains limited to a few empirical observations [35, 40-43].

In this work, we conducted an exhaustive analysis of the GNRA tetraloop motif instances across known 3D structures of nucleic acids, without imposing any restraints on sequence, loop type, backbone topology, or involved interactions. We observed four distinct insertion sites in the pentaloop variants of the motif, with three of them forming interactions with proteins. We identified twelve recurrent backbone topology variants of the GNRA structure, including all five possible two-strand variants and six out of ten theoretically possible three-strand variants. These GNRA-like motifs illustrate the remarkable capacity of favorable RNA 3D motifs to fit in diverse backbone contexts, adding another dimension to the complexity of RNA folding.

## RESULTS

We defined the GNRA tetraloop motif as consisting of six residues (n1 to n6), which include the GNRA tetraloop (n2 to n5) and one flanking canonical base pair n1-n6 (Figure 1). To explore the landscape of GNRA-like motif variants, we selected the reference instance (Figure 1A) and searched for its matches in nucleic acid-containing PDB entries [44] using the ARTEM tool [42], see the Methods section for details.

### Dataset overview

We identified 23,283 non-redundant GNRA tetraloop matches comprising three to six residues across 3,122 PDB entries (Supplementary Table S1), representing 13,729 structurally unique motifs (Supplementary Table S2). These matches were found in 86 non-coding RNA families from Rfam [45] and 1,063 classes from the BGSU representative set of RNA structures [46], with only 22 matches involving DNA residues. The list of Rfam families included 21 riboswitch aptamers, 19 ribozymes, 19 viral RNAs, including 4 IRES elements, 9 spliceosomal RNAs, and 8 ribosomal RNAs. Among the most frequently observed modifications were PSU (199 matches), A2M (162), MA6 (133), OMG (104), and OMC (97). 10,894 hits matched all six residues of the reference. The n1 and n6 positions were identified as the least constrained, with 4,068 and 6,061 unmatched cases, respectively, followed by n5 (3,038) and n3 (2,879). Notably, among the n2 and n4 residues, both critical for the GNRA tetraloop motif and the U-turn motif, the n2 was found unmatched nearly five times more often than n4 (2,393 vs. 521 cases, respectively).

The n2 and n5 residues were identified as the most conserved positions, with 72% and 78% of the matched bases being G and A, respectively (Figure 2). The n4 position matched a purine in 80% of cases, and, notably, adenosines accounted for 57% of the matched n3 residues. 4,881 hits with matched n2-n5 residues were formed by tetraloops, 1,409 by pentaloops, 809 by hexaloops, and 2,303 by longer hairpin loops. The remaining 13,881 matches either included non-hairpin strands or had at least one unmatched residue from n2 to n5.

**Figure 2.**
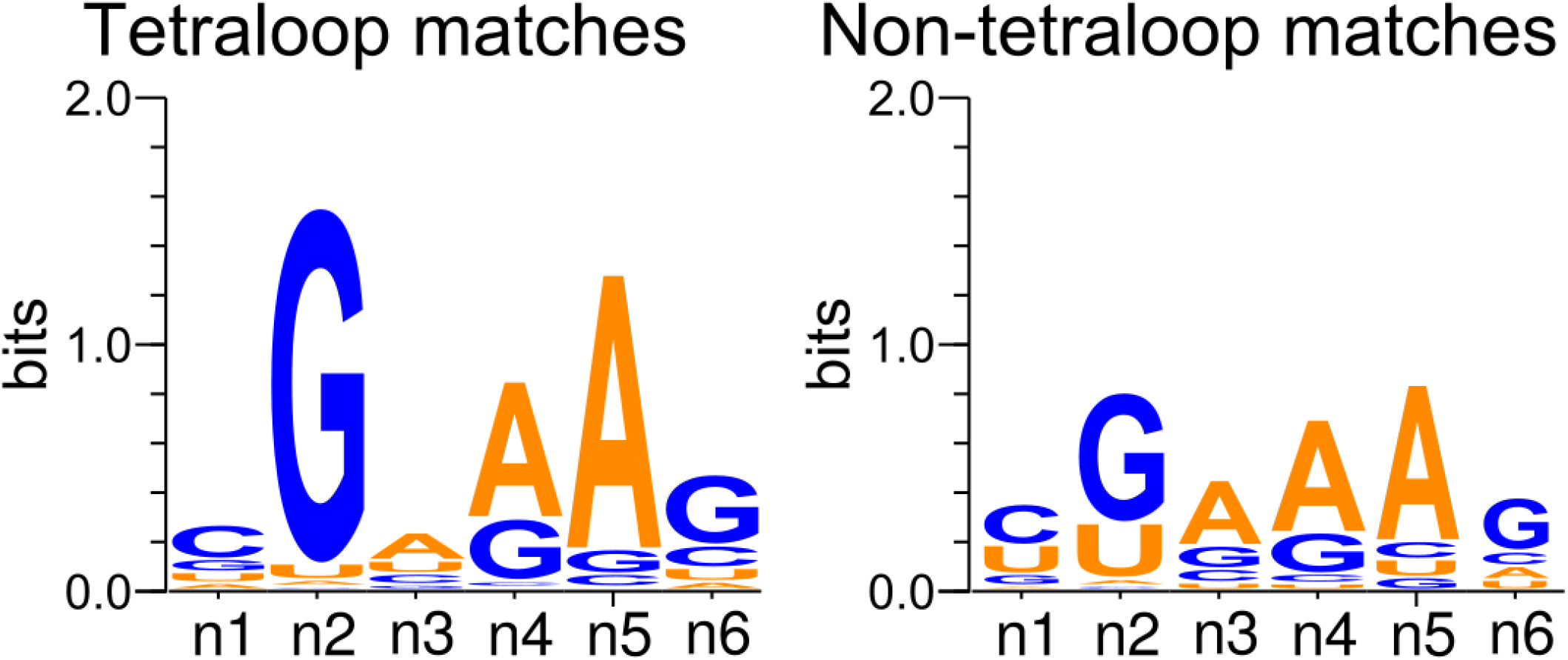
Motif logo of tetraloop and non-tetraloop matches of the GNRA tetraloop motif. The left logo shows the 4,881 tetraloop matches with a single continuous n1-n6 strand. The right logo shows the remaining 18,402 matches. The figure was prepared using WebLogo 3 (https://weblogo.threeplusone.com/create.cgi).

ARTEM successfully captured known variants of the GNRA tetraloop motif. Among others, the dataset included the DNA form of the motif from a gapped dumbbell DNA [47], which retained all characteristic interactions except the n2-n4 R-rG base-ribose hydrogen bond (Figure 3A); the UAA/GAN internal loop, which lacked the n2-n5 G-RpA base-phosphate interaction and the n2-n4 NpR-G oxygen-base stacking, instead forming an n2-n4 G-NpR base-phosphate hydrogen bond (Figure 3B); and the UMAC structure formed by the CUAACC hexaloop of the bacteriophage φ21 boxB RNA in complex with the N peptide [27], which lacked the n2-n5 tSH base pair and featured a non-canonical cWW C-C base pair as the flanking n1-n6 interaction (Figure 3C).

**Figure 3.**
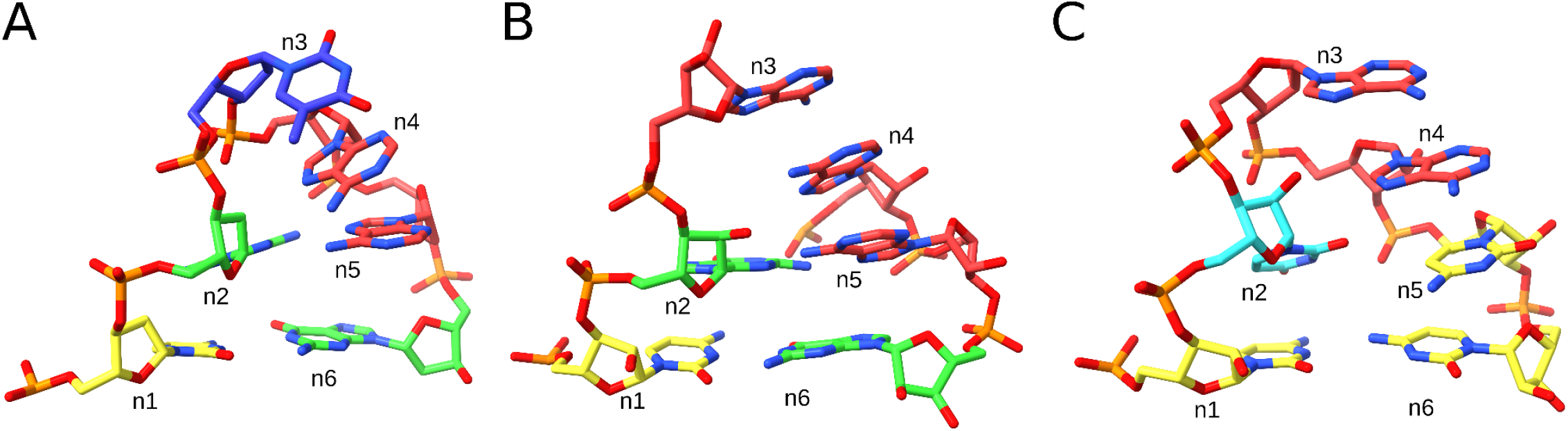
Known GNRA-like motif variants captured by ARTEM. (**A**) DNA form, GTAA tetraloop, 1.04 Å RMSD, PDB entry 2N8A, chain B, residues 9-14. (**B**) UAA/GAN internal loop, 1.25 Å RMSD, PDB entry 4V72, 23S rRNA, chain BA, residues 1376-1378 and 1353-1355. (**C**) CUAACC hexaloop, 1.47 Å RMSD, PDB entry 1NYB, phage φ21 boxB RNA, chain B, residues 10-15. The color scheme shows guanosines in green, adenosines in red, cytidines in yellow, and thymidines and uridines in blue. The figure was prepared using UCSF ChimeraX (https://www.cgl.ucsf.edu/chimerax/).

Among the 13,729 unique motifs, 236 were identified in at least three different Rfam families and three different BGSU representative classes and can therefore be considered recurrent. The GAAA tetraloop, which did not form long-range base pairs, was found in eight Rfam families with a G-C flanking n1-n6 base pair, including group II introns, SAM and Glutamine riboswitches, and bacterial RNase P, and in seven families with a C-G flanking base pair, including aptamers of Lysine, Guanidine-I, and AdoCbl-II riboswitches. The GUGA tetraloop, in which the n4 and n5 residues formed long-range base pairs, was identified in six Rfam families with a G-C flanking n1-n6 base pair, including the glmS ribozyme and ribosomal RNAs. All other motifs were found in at most five Rfam families, with a UAA/GAN six-residue match, identified in 21 representative classes across three families of ribosomal RNAs, being the most widespread non-hairpin motif.

### Pentaloop variants

We analyzed the identified pentaloop variants of the GNRA tetraloop motif. With ARTEM, we did not detect any pentaloop variants with an insertion between n1 and n2 but identified all four of the other possible insertion variants (Figure 4). We selected non-ribosomal matches with the lowest RMSD to the reference as representative instances. The n2-n3 insertion instance (Figure 4A, B) was found in a viral tRNA-like structure [48], with an extra uracil stacked on top of n3, and did not form any external interactions. The three remaining representatives interacted with proteins.

**Figure 4.**
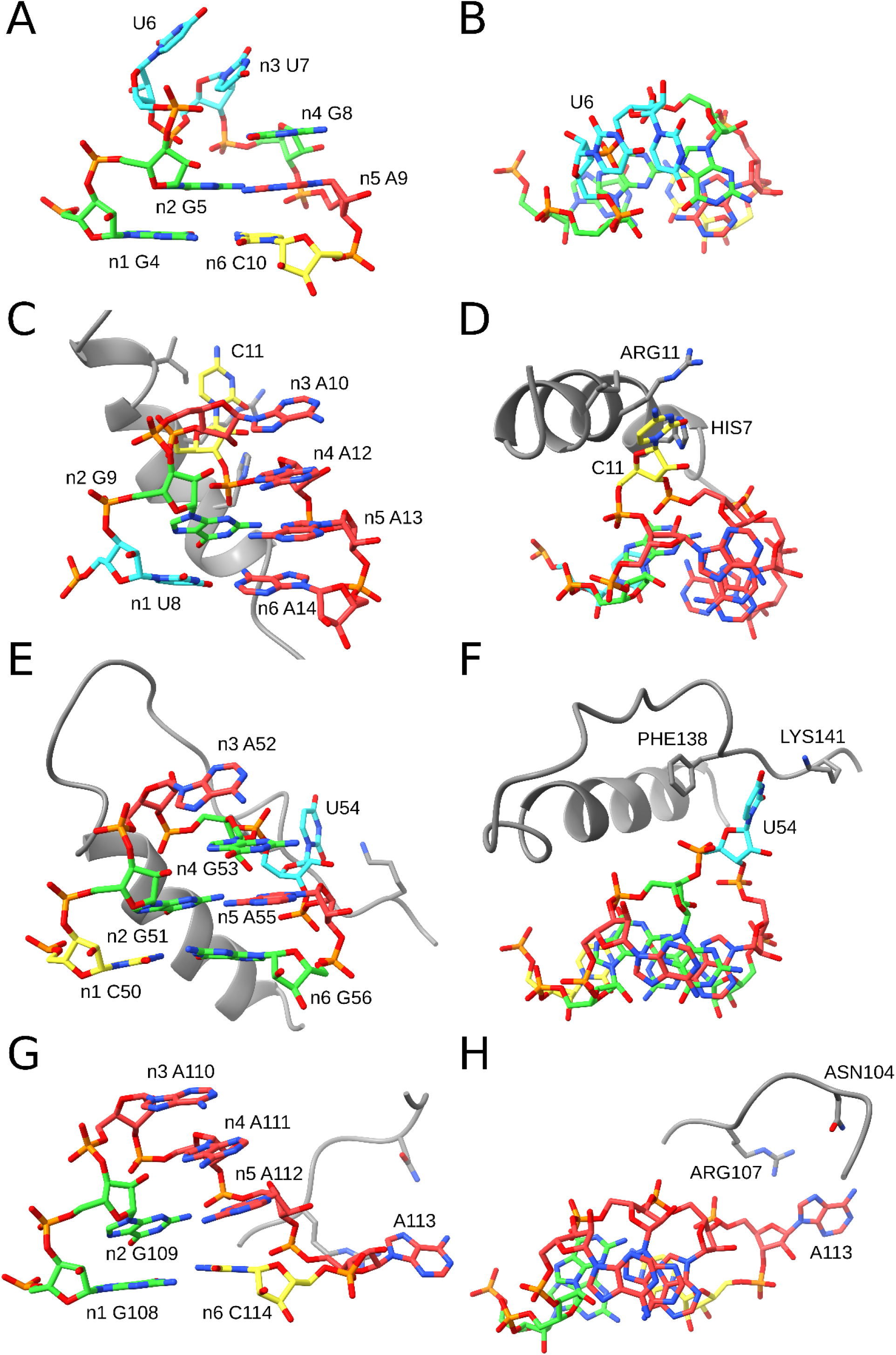
Pentaloop variants of the GNRA tetraloop motif captured by ARTEM. The left column is a side view, and the right column is a top view. (**A, B**) n2-n3 insertion variant from a viral tRNA-like structure, 1.28 Å RMSD, PDB entry 7SC6, chain C. (**C, D**) n3-n4 insertion variant from the phage P22 N-peptide-BoxB RNA complex, 1.15 Å RMSD, PDB entry 1A4T, chain A. (**E, F**) n4-n5 insertion variant from a CRISPR RNA, 0.84 Å RMSD, PDB entry 6LNB, chain M. (**G, H**) n5-n6 insertion variant from an RNase MRP structure, 1.03 Å RMSD, PDB entry 6W6V, chain A. The color scheme shows guanosines in green, adenosines in red, cytidines in yellow, and thymidines and uridines in blue. The interacting proteins are shown in gray. The figure was prepared using UCSF ChimeraX (https://www.cgl.ucsf.edu/chimerax/).

The n3-n4 insertion variant was found in the N-peptide-BoxB RNA complex of bacteriophage P22 [24] (Figure 4C, D), with a cytosine looped out from the GACAA pentaloop and interacting with arginine ARG11 (O2-NE distance of 3.3 Å) and histidine HIS7 (O2-NE2 distance of 3.6 Å) of the N-peptide. The n4-n5 insertion variant was found in a CRISPR RNA structure [49] (Figure 4E, F). The U54 residue was looped out and formed a stacking interaction with phenylalanine PHE138 (∼5 Å distance between the ring planes) and a hydrogen bond with lysine LYS141 (O2-NZ distance of 4.2 Å) of the Cas6 protein. The n5-n6 insertion variant was observed in an RNase MRP structure [50] (Figure 4G, H). The looped out adenosine A113 was exposed to arginine ARG107 (N7-NH2 distance of 4.5 Å) and asparagine ASN104 (N6-NO2 distance of 5.7 Å) of the ribonucleases P/MRP protein subunit POP1.

### Backbone topology landscape of GNRA-like motif variants

We classified all 13,729 motifs into 59 distinct backbone topologies (Supplementary Table S3), based on binary relationships between neighboring positions in the GNRA tetraloop motif. For example, in the canonical GNRA tetraloop, all neighboring positions are consecutive residues, yielding the backbone topology “n1-n2-n3-n4-n5-n6”. In contrast, in the UAA/GAN motif, positions n3 and n4 are not consecutive in the RNA sequence, resulting in the “n1-n2-n3|n4-n5-n6” topology. In total, we identified 17 distinct topologies among the six-residue matches, out of the 2^5^ = 32 possible variants; 12 of these were found in at least three BGSU classes and three Rfam families (Figure 5, Table 1). We then analyzed representative motifs from each variant, selecting those with the lowest RMSD to the reference. When a representative motif contained artifacts, for example, overlapping residues in low-resolution entries, we selected an appropriate next-best suitable match.

**Table 1.**
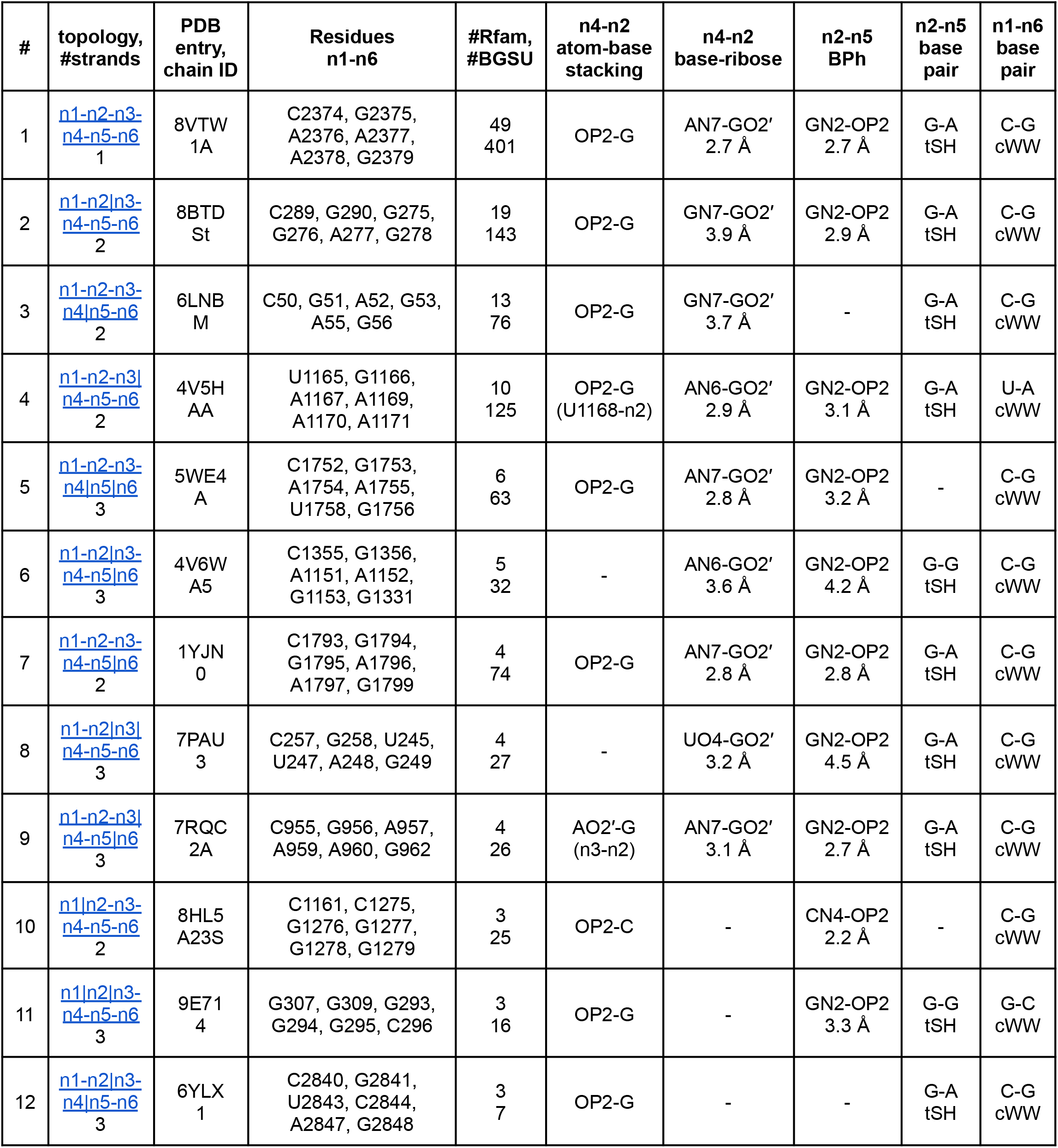
Representative motifs of the recurrent backbone topology variants of the GNRA tetraloop motif.

**Figure 5.**
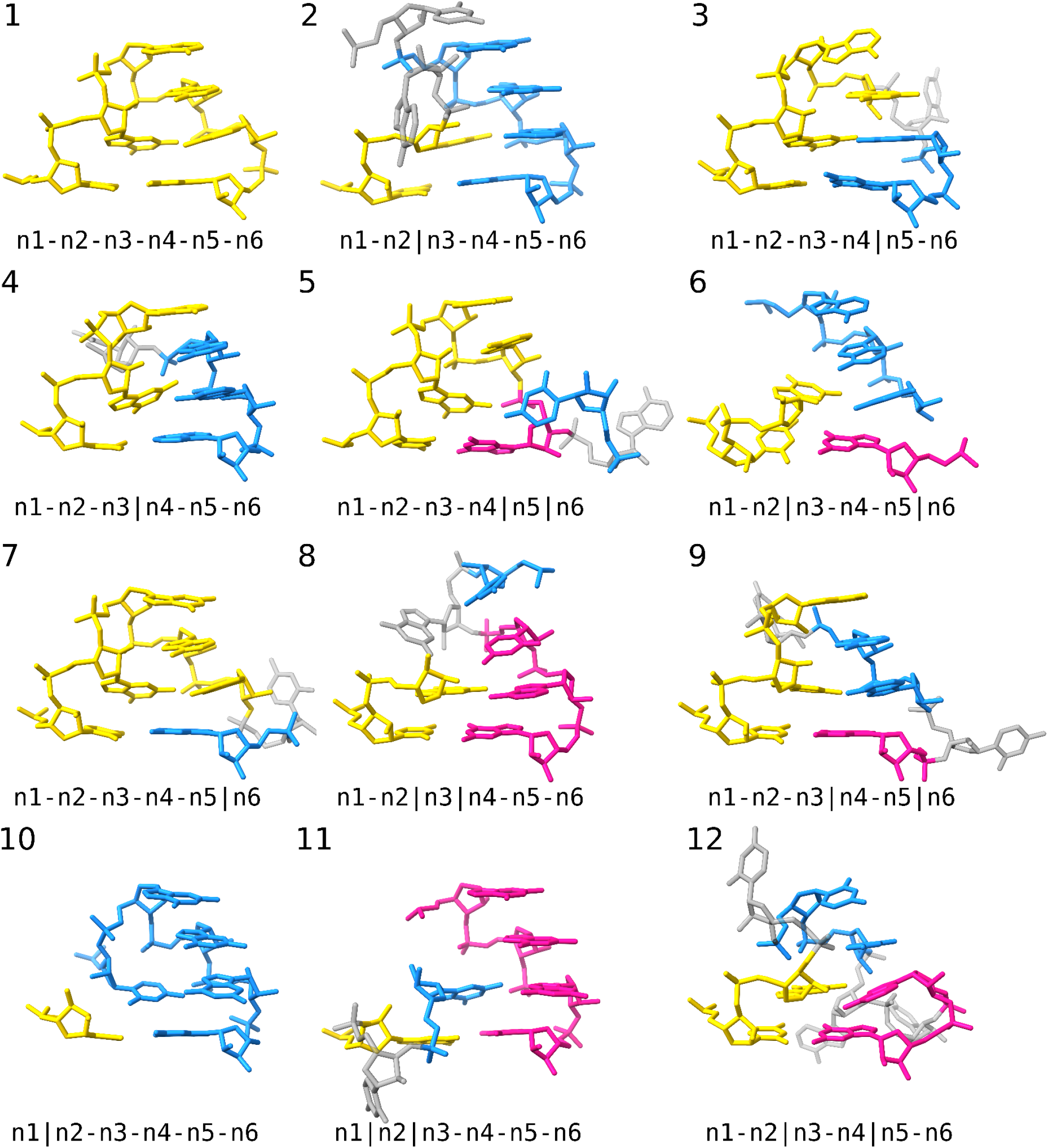
Recurrent backbone topology variants of the GNRA tetraloop motif identified by ARTEM. Continuous strands of the motif are shown in gold, blue, and pink. Looped-out residues are shown in gray. The figure was prepared using UCSF ChimeraX (https://www.cgl.ucsf.edu/chimerax/).

As expected, the canonical one-strand variant of the GNRA tetraloop motif was the most frequent motif, identified in 49 different Rfam families. It was followed by three two-strand variants with a single chain break: the n2|n3 break found in 19 Rfam families, the n4|n5 break in 13 families, and the n3|n4 variant in 10 families. The remaining two-strand variants were less common, with n5|n6 found in four (rRNAs and RNase MRP) and n1|n2 found in three Rfam families (rRNAs only). The two most frequent three-strand variants were “n1-n2-n3-n4|n5|n6” and “n1-n2|n3-n4-n5|n6”, identified in six (rRNAs, group II introns, and FMN riboswitch) and five Rfam families (rRNAs only), respectively. Representative motifs of all twelve topologies featured at least three of the five key interactions characteristic of the GNRA tetraloop motif (Table 1).

We surveyed the most populated three-strand “n1-n2-n3-n4|n5|n6” topology to assess the diversity of instances in terms of key interactions (Figure 6, Table 2). In addition to the representative motif from 23S rRNA, which lacks an n2-n5 base pair (Figure 6A), we identified instances featuring n2-n5 base pairs involving each of the three possible edges of the n5 residue. A match from a group II intron included U-A cSW (cis-Sugar-Watson-Crick) n2-n5 base pair and the canonical U-A n1-n6 base pair (Figure 6B); a 16S rRNA match featured a G-U cSS n2-n5 base pair (Figure 6C); and another 16S rRNA match displayed the classic G-A tSH n2-n5 base pair along with a non-canonical U-A tWH n1-n6 base pair (Figure 6D). All four instances contained the other three key interactions characteristic of the GNRA tetraloop motif (Table 2).

**Table 2.**
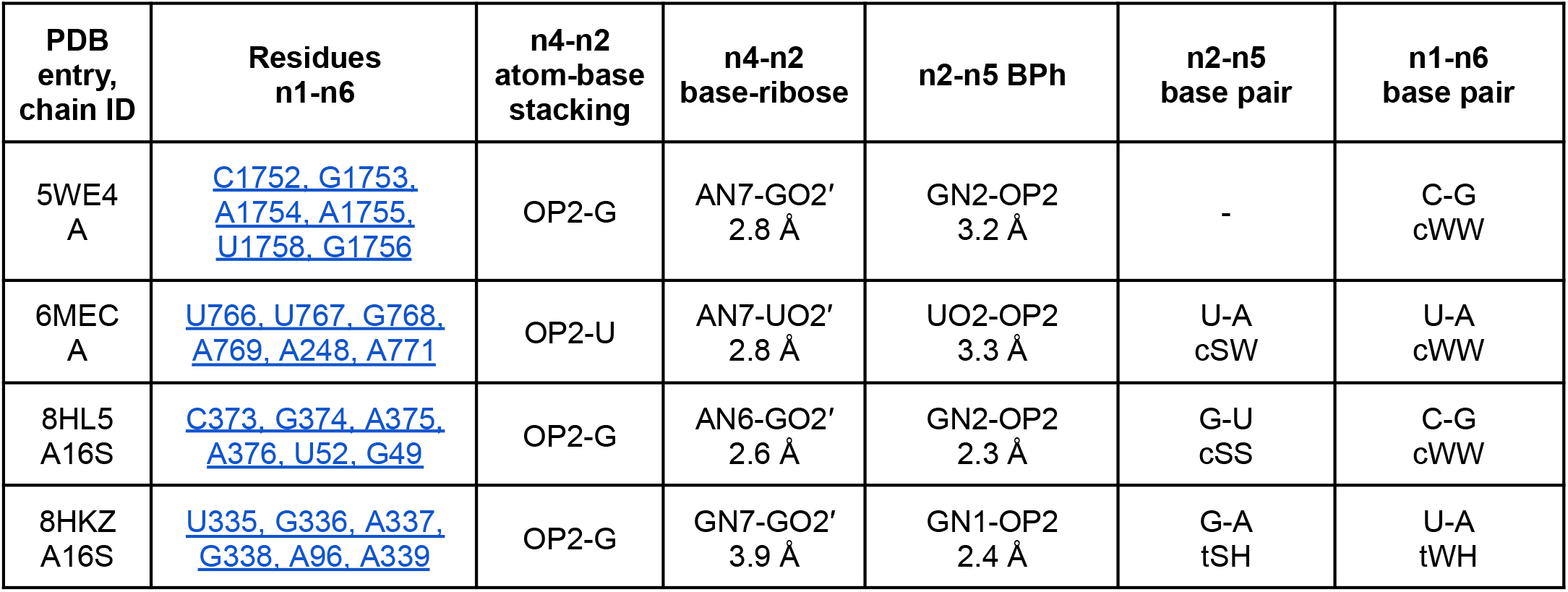
Four representative motifs of the GNRA-like topology variant n1-n2-n3-n4|n5|n6 (#5).

**Figure 6.**
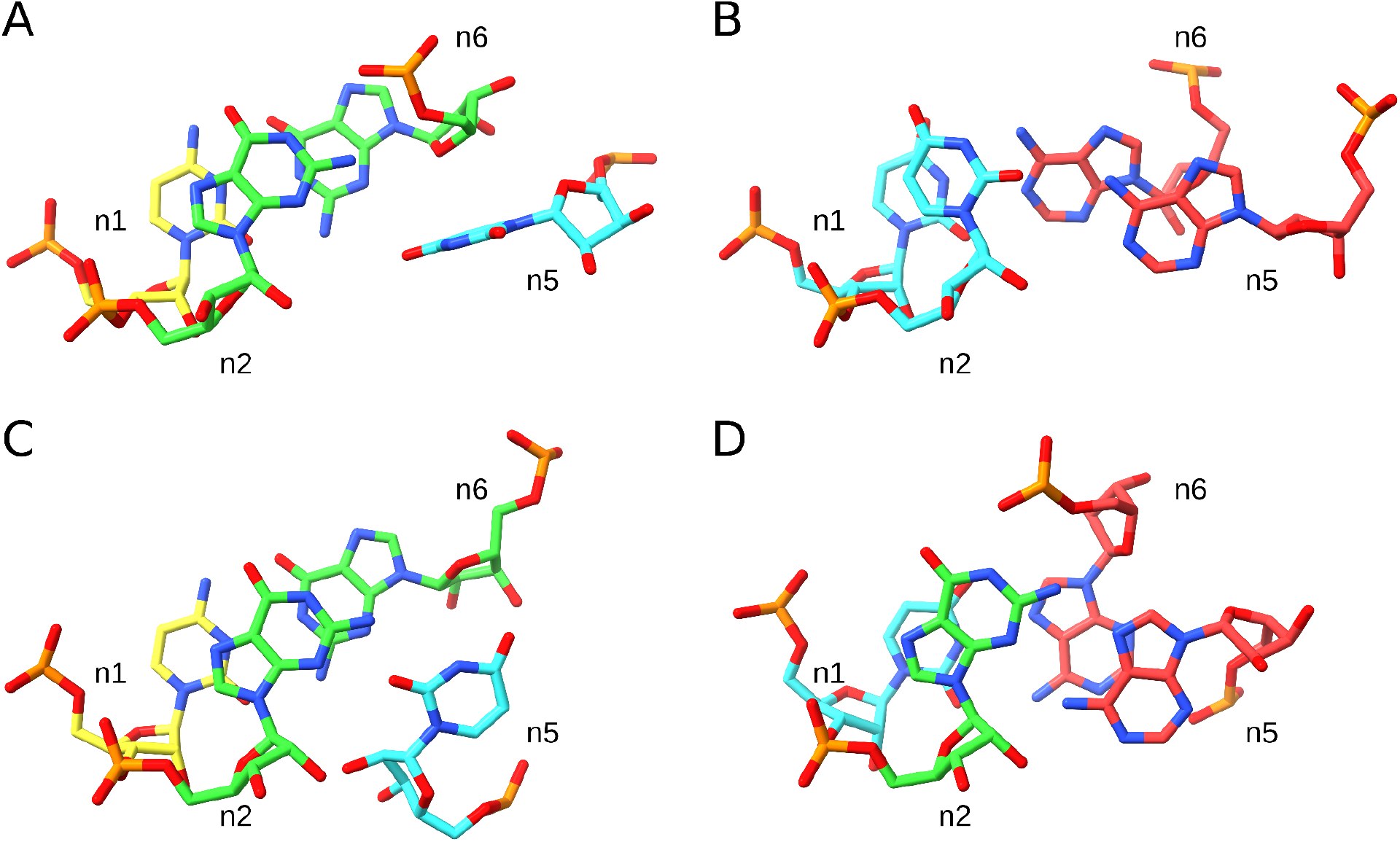
Base pair diversity of the GNRA-like topology variant n1-n2-n3-n4|n5|n6 (#5). (**A**) 23S rRNA, 0.91 Å RMSD match, PDB entry 5WE4, chain A. (**B**) Group II intron, 1.23 Å RMSD match, PDB entry 6MEC, chain A. (**C**) 16S rRNA, 1.44 Å RMSD match, PDB entry 8HL5, chain A16S. (**D**) 16S rRNA, 1.49 Å RMSD match, PDB entry 8HKZ, chain A16S. The color scheme shows guanosines in green, adenosines in red, cytidines in yellow, and uridines in blue. The figure was prepared using UCSF ChimeraX (https://www.cgl.ucsf.edu/chimerax/).

## DISCUSSION

In this work, we conducted a comprehensive survey of GNRA tetraloop-like motifs without setting any restraints on sequence, interactions, or backbone topology. In addition to previously known non-canonical variants of the motif, including DNA forms, UMAC sequences, and UAA/GAN internal loops, we identified several new GNRA-like backbone topology variants. These included all possible two-strand variants, with n2|n3, n3|n4, and n4|n5 chain breaks among the most frequent, and n1|n2 and n5|n6 breaks less favored. We also identified six three-strand variants of the GNRA-like motif, including the recurrent “n1-n2-n3-n4|n5|n6” topology, which accommodates a diverse range of n2-n5 base pair geometries. Furthermore, we observed four different insertion locations in the pentaloop variants of the GNRA tetraloop motif, three of which were involved in RNA-protein interactions.

The fact that the n1|n2 chain break was identified as the least frequent two-strand variant of the GNRA tetraloop motif, and was not found among the pentaloop variants, aligns with the previously noted importance of n1-n2 connectivity in forming the motif [30]. However, the diversity of identified GNRA-like backbone topology variants suggests that the backbone acts as a contributing factor rather than a strict determinant. Overall, no definitive determinants of the GNRA structure have been identified, as it can accommodate non-GNRA sequences, one to three strands, RNA or DNA residues, and various types of core interactions. Examining the representatives of the identified GNRA-like variants (Figure 5, Table 1), it is not obvious where to place a clear demarcation to decide what does and does not qualify as a GNRA tetraloop-like motif. In this work, the principle of the ARTEM tool, to identify matches based on motif isostericity (the 3D alignability of residues), served as the only constraint enabling such separation. In contrast, defining the GNRA tetraloop motif based on the centroid reference (Figure 1) is straightforward. Thus, we observe that the GNRA tetraloop motif, like arbitrary RNA tertiary motifs in general, can be viewed as a clearly defined, energetically favorable gravity center surrounded by a fuzzy cloud of less favorable variants. It is important to note, however, that we do not account for the structural environment of the motif and therefore cannot distinguish between self-sufficient instances and those that depend on specific interaction partners. The ability to differentiate between these cases could, in principle, provide the desired criterion for classification.

The diversity of RNA-protein interaction modes observed among the pentaloops captured by ARTEM highlights the biological relevance of the GNRA tetraloop motif variants. In the analyzed representative pentaloop instances, the looped-out bases were directly exposed to proteins, contributing to the specificity of the intermolecular interactions. Thus, systematic analysis of RNA tertiary motif variability, such as the investigation of GNRA tetraloop motifs presented in this work, serves as a step toward decoding the RNA-RNA, RNA-protein, and RNA-ligand recognition modes and holds promise for modulating binding affinities through RNA motif design.

While ARTEM proves to be a powerful tool for the analysis of RNA 3D motifs, it has two notable limitations. First, it requires the user to define the RNA motif of interest. Second, its results are biased toward the selected reference motif instances. In future work, we plan to equip ARTEM with a built-in library of recurrent RNA 3D motifs, which will significantly expand the tool’s applicability. Ultimately, providing the research community with a comprehensive resource will make nucleic acid 3D structural data more interpretable and actionable for non-structural biologists.

## METHODS

To survey GNRA-like motifs, we retrieved all PDB entries containing RNA, DNA, or hybrid nucleic acid molecules (18,741 entries as of January 13th, 2025, mmCIF format [44]). We retained only the first model in each entry and added interacting symmetry mates using ChimeraX [51] (“*crystalcontacts #1 distance 6*”) to account for crystal contacts (only for entries under 10MB with the experimental method “X-RAY DIFFRACTION”). To select the reference GNRA tetraloop instance (23S rRNA, entry 8VTW, chain 1A, residues 2374-2379) [52], we identified the largest hairpin loop motif annotated as “GNRA” in the RNA 3D Motif Atlas [53] (motif HL_85603.2, Hairpin Loop Motif Atlas Release 3.88), and searched for the centroid instance, defined as the one with the lowest median RMSD to other instances of the motif, using ARTEM [42] for pairwise comparisons.

We then searched for matches of the GNRA reference in the PDB entries using ARTEM in two modes: (i) using the six-residue reference (n1 to n6, GNRA plus the flanking base pair), counting matches of five residues with an RMSD under 1.25 Å and six residues with an RMSD under 1.5 Å; (ii) using the four-residue reference (n2 to n5, GNRA only), counting matches of three residues with an RMSD under 0.75 Å that include the n2 residue and matches of four residues with an RMSD under 1.0 Å. The RMSD thresholds were defined based on pairwise RMSD distributions between instances of the HL_85603.2 motif (Supplementary Figure S1). The two-mode approach was designed to avoid numerous trivial two-base-pair hits (the G-A base pair plus the flanking base pair) that would be identified by ARTEM if searched with the six-residue reference and counting four-residue matches. Each search took less than eight hours on an Asus VivoBook 15 laptop. The matches obtained from the two runs were merged, retaining only those smaller matches that were not subsets of any larger match. This process resulted in 133,560 hits. Each residue of a hit was assigned its reference counterpart, from n1 (reference residue 2374) to n6 (reference residue 2379).

The hits were then annotated with structural features using urslib2 [42], DSSR (version 2.0, [54]), and custom Python scripts. This included per-residue crystal contacts (non-hydrogen atoms from symmetry mates within 4.0 Å), base types of the matched residues, sequence distance between consecutive residues of the match (e.g., 1 for direct neighbors), RNA secondary structure elements (stems and loops) for each residue, and their relationships between consecutive residues (same element, local, long-range), hairpin length and sequence (for hairpin matches), base orientations (anti/syn) and sugar pucker conformations, n2-n5 base pair type according to the Leontis-Westhof (LW) nomenclature [55], the LW type of the flanking n1-n6 base pair, per-residue information on base-pairing and base-staking interactions with other residues outside the match, U-turn formation, and Rfam families [45] and BGSU representative set classes (version 3.370, [46]) of the involved RNA chains.

To account for redundancy, we designed a descriptor string for each hit, which included all the annotated structural features, Rfam families, and BGSU representative set classes. We then merged hits with identical descriptors, keeping the lowest-RMSD hit as the representative. This resulted in a non-redundant set of 23,283 instances (Supplementary Table S1). Subsequently, we merged the instances using descriptors based only on structural features, counting the different Rfam families and BGSU representative set classes annotated for each descriptor, and retaining the lowest-RMSD hit as representative. This produced 13,729 structurally unique hits (Supplementary Table S2). Similarly, we obtained a set of 59 representatives of unique strand topologies (Supplementary Table S3). The use of RNA-focused annotations (Rfam family, BGSU representative set class) was supported by the fact that only 22 of the 23,283 non-redundant hits contained DNA bases (DA, DC, DG, or DT).

A step-by-step description of the performed work is available in the GitHub repository (https://github.com/febos/GNRA/blob/main/REPRODUCE.md).

## DATA AVAILABILITY

The code and data are available at https://github.com/febos/GNRA (DOI: https://doi.org/10.5281/zenodo.15670527).

## ACKNOWLEDGMENTS

This work was supported by the National Science Center Poland (NCN) [grant 2017/25/B/NZ2/01294 to J.M.B., grant 2024/53/B/NZ2/02718 to E.F.B.].

## SUPPLEMENTARY MATERIALS

**Supplementary Figure S1.**
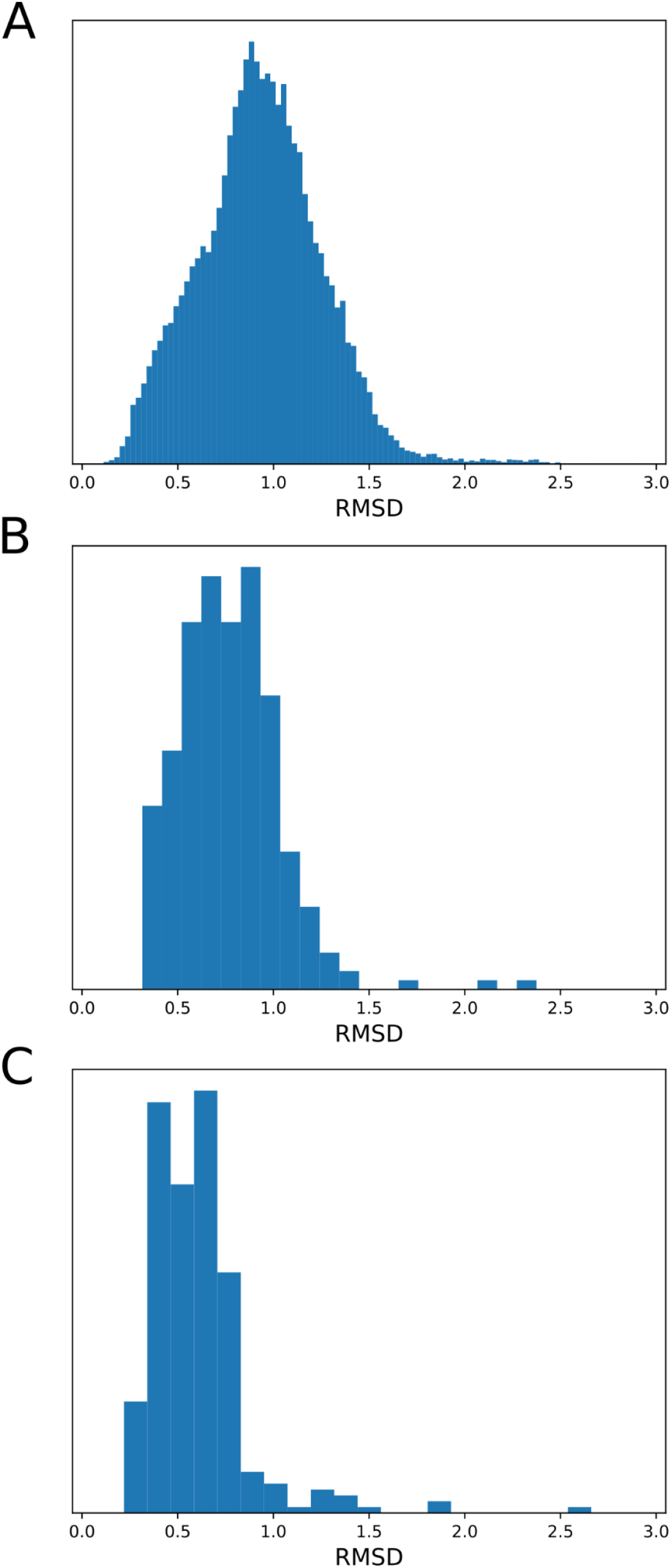
Pairwise RMSD values among the 284 GNRA motif instances of the HL_85603.2 motif reported by ARTEM for (**A**) six-residue matches (GNRA loop and flanking base pair) among all possible pairs, (**B**) six-residue matches with respect to the centroid instance (23S rRNA, PDB entry 8VTW, chain 1A, residues 2374-2379), and (**C**) four-residue matches with the centroid instance (GAAA loop, residues 2375-2378).

## REFERENCES

1. Zhang, H., & Ding, Y. (2025). RNA Structure: Function and Application in Plant Biology. Annual Review of Plant Biology, 76. 10.1146/annurev-arplant-083123-055521

2. Ganser, L. R., Kelly, M. L., Herschlag, D., & Al-Hashimi, H. M. (2019). The roles of structural dynamics in the cellular functions of RNAs. Nature reviews Molecular cell biology, 20(8), 474–489. 10.1038/s41580-019-0136-0

3. Travers, A., & Muskhelishvili, G. (2015). DNA structure and function. The FEBS journal, 282(12), 2279–2295. 10.1111/febs.13307

4. Collie, G. W., & Parkinson, G. N. (2011). The application of DNA and RNA G-quadruplexes to therapeutic medicines. Chemical Society Reviews, 40(12), 5867–5892. 10.1039/C1CS15067G

5. Ohno, H., Akamine, S., & Saito, H. (2019). RNA nanostructures and scaffolds for biotechnology applications. Current Opinion in Biotechnology, 58, 53–61. 10.1016/j.copbio.2018.11.006

6. Stombaugh, J., Zirbel, C. L., Westhof, E., & Leontis, N. B. (2009). Frequency and isostericity of RNA base pairs. Nucleic acids research, 37(7), 2294–2312. 10.1093/nar/gkp011

7. Butcher, S. E., & Pyle, A. M. (2011). The molecular interactions that stabilize RNA tertiary structure: RNA motifs, patterns, and networks. Accounts of chemical research, 44(12), 1302–1311. 10.1021/ar200098t

8. Uhlenbeck, O. C. (1990). Tetraloops and RNA folding. Nature, 346(6285), 613–614. 10.1038/346613a0

9. Woese, C. R., Winker, S., & Gutell, R. R. (1990). Architecture of ribosomal RNA: constraints on the sequence of” tetra-loops”. Proceedings of the National Academy of Sciences, 87(21), 8467–8471. 10.1073/pnas.87.21.8467

10. Antao, V. P., Lai, S. Y., & Tinoco Jr, I. (1991). A thermodynamic study of unusually stable RNA and DNA hairpins. Nucleic acids research, 19(21), 5901–5905. 10.1093/nar/19.21.5901

11. Heus, H. A., & Pardi, A. (1991). Structural features that give rise to the unusual stability of RNA hairpins containing GNRA loops. Science, 253(5016), 191–194. 10.1126/science.1712983

12. Jucker, F. M., & Pardi, A. (1995). GNRA tetraloops make a U-turn. Rna, 1(2), 219. https://pmc.ncbi.nlm.nih.gov/articles/PMC1369075/

13. d’Ascenzo, L., Leonarski, F., Vicens, Q., & Auffinger, P. (2017). Revisiting GNRA and UNCG folds: U-turns versus Z-turns in RNA hairpin loops. Rna, 23(3), 259–269. 10.1261/rna.059097.116

14. Michel, F., & Westhof, E. (1990). Modelling of the three-dimensional architecture of group I catalytic introns based on comparative sequence analysis. Journal of molecular biology, 216(3), 585–610. 10.1016/0022-2836(90)90386-Z

15. Jaeger, L., Michel, F., & Westhof, E. (1994). Involvement of a GNRA tetraloop in long-range RNA tertiary interactions. Journal of molecular biology, 236(5), 1271–1276. 10.1016/0022-2836(94)90055-8

16. Geary, C., Baudrey, S., & Jaeger, L. (2008). Comprehensive features of natural and in vitro selected GNRA tetraloop-binding receptors. Nucleic acids research, 36(4), 1138–1152. 10.1093/nar/gkm1048

17. Wu, L., Chai, D., Fraser, M. E., & Zimmerly, S. (2012). Structural variation and uniformity among tetraloop-receptor interactions and other loop-helix interactions in RNA crystal structures. PLoS One, 7(11), e49225. 10.1371/journal.pone.0049225

18. Fiore, J. L., & Nesbitt, D. J. (2013). An RNA folding motif: GNRA tetraloop–receptor interactions. Quarterly reviews of biophysics, 46(3), 223–264. 10.1017/S0033583513000048

19. Fernández-Miragall, O., & Martínez-Salas, E. (2003). Structural organization of a viral IRES depends on the integrity of the GNRA motif. Rna, 9(11), 1333–1344. 10.1261/rna.5950603

20. Robertson, M. E., Seamons, R. A., & Belsham, G. J. (1999). A selection system for functional internal ribosome entry site (IRES) elements: analysis of the requirement for a conserved GNRA tetraloop in the encephalomyocarditis virus IRES. Rna, 5(9), 1167–1179. 10.1017/S1355838299990301

21. Psaridi, L., Georgopoulou, U., Varaklioti, A., & Mavromara, P. (1999). Mutational analysis of a conserved tetraloop in the 5′ untranslated region of hepatitis C virus identifies a novel RNA element essential for the internal ribosome entry site function. FEBS letters, 453(1-2), 49–53. 10.1016/S0014-5793(99)00662-6

22. Correll, C. C., Munishkin, A., Chan, Y. L., Ren, Z., Wool, I. G., & Steitz, T. A. (1998). Crystal structure of the ribosomal RNA domain essential for binding elongation factors. Proceedings of the National Academy of Sciences, 95(23), 13436–13441. 10.1073/pnas.95.23.13436

23. García-Ortega, L., Álvarez-García, E., Gavilanes, J. G., Martínez-del-Pozo, Á., & Joseph, S. (2010). Cleavage of the sarcin–ricin loop of 23S rRNA differentially affects EF-G and EF-Tu binding. Nucleic acids research, 38(12), 4108–4119. 10.1093/nar/gkq151

24. Cai, Z., Gorin, A., Frederick, R., Ye, X., Hu, W., Majumdar, A., … & Patel, D. J. (1998). Solution structure of P22 transcriptional antitermination N peptide–box B RNA complex. Nature structural biology, 5(3), 203–212. 10.1038/nsb0398-203

25. Schärpf, M., Sticht, H., Schweimer, K., Boehm, M., Hoffmann, S., & Rösch, P. (2000). Antitermination in bacteriophage λ: The structure of the N36 peptide-boxB RNA complex. European Journal of Biochemistry, 267(8), 2397–2408. 10.1046/j.1432-1327.2000.01251.x

26. Saon, M. S., Kirkpatrick, C. C., & Znosko, B. M. (2023). Identification and characterization of RNA pentaloop sequence families. NAR genomics and bioinformatics, 5(1), lqac102. 10.1093/nargab/lqac102

27. Cilley, C. D., & Williamson, J. R. (2003). Structural mimicry in the phage [phis] 21 N peptide–boxB RNA complex. Rna, 9(6), 663–676. 10.1261/rna.2189203

28. Cheong, C., & Cheong, H. K. (2010). RNA structure: tetraloops. eLS. 10.1002/9780470015902.a0003135.pub2

29. Thapar, R., Denmon, A. P., & Nikonowicz, E. P. (2014). Recognition modes of RNA tetraloops and tetraloop-like motifs by RNA-binding proteins. Wiley Interdisciplinary Reviews: RNA, 5(1), 49–67. 10.1002/wrna.1196

30. Abramovitz, D. L., & Pyle, A. M. (1997). Remarkable morphological variability of a common RNA folding motif: the GNRATetraloop-receptor interaction. Journal of molecular biology, 266(3), 493–506. 10.1006/jmbi.1996.0810

31. Moody, E. M., Feerrar, J. C., & Bevilacqua, P. C. (2004). Evidence that folding of an RNA tetraloop hairpin is less cooperative than its DNA counterpart. Biochemistry, 43(25), 7992–7998. 10.1021/bi049350e

32. Lee, J. C., Gutell, R. R., & Russell, R. (2006). The UAA/GAN internal loop motif: a new RNA structural element that forms a cross-strand AAA stack and long-range tertiary interactions. Journal of molecular biology, 360(5), 978–988. 10.1016/j.jmb.2006.05.066

33. Jaeger, L., Verzemnieks, E. J., & Geary, C. (2009). The UA_handle: a versatile submotif in stable RNA architectures. Nucleic acids research, 37(1), 215–230. 10.1093/nar/gkn911

34. Geary, C., Chworos, A., & Jaeger, L. (2011). Promoting RNA helical stacking via A-minor junctions. Nucleic acids research, 39(3), 1066–1080. 10.1093/nar/gkq748

35. Jiang, F., Kumar, R. A., Jones, R. A., & Patel, D. J. (1996). Structural basis of RNA folding and recognition in an AMP–RNA aptamer complex. Nature, 382(6587), 183–186. 10.1038/382183a0

36. Huang, H. C., Nagaswamy, U. M. A., & Fox, G. E. (2005). The application of cluster analysis in the intercomparison of loop structures in RNA. Rna, 11(4), 412–423. 10.1261/rna.7104605

37. Baulin, E. F., Bohdan, D. R., Kowalski, D., Serwatka, M., Świerczyńska, J., Żyra, Z., & Bujnicki, J. M. (2024). ARTEM: a method for RNA and DNA tertiary motif identification with backbone permutations, and its example application to kink-turn-like motifs. bioRxiv, 2024-05. 10.1101/2024.05.31.596898

38. Zhao, Q., Huang, H. C., Nagaswamy, U., Xia, Y., Gao, X., & Fox, G. E. (2012). UNAC tetraloops: to what extent do they mimic GNRA tetraloops?. Biopolymers, 97(8), 617–628. 10.1002/bip.22049

39. Bottaro, S., & Lindorff-Larsen, K. (2017). Mapping the universe of RNA tetraloop folds. Biophysical journal, 113(2), 257–267. 10.1016/j.bpj.2017.06.011

40. Trachman III, R. J., Demeshkina, N. A., Lau, M. W., Panchapakesan, S. S. S., Jeng, S. C., Unrau, P. J., & Ferré-D’Amaré, A. R. (2017). Structural basis for high-affinity fluorophore binding and activation by RNA Mango. Nature chemical biology, 13(7), 807–813. 10.1038/nchembio.2392

41. Liu, Y., Wilson, T. J., & Lilley, D. M. (2017). The structure of a nucleolytic ribozyme that employs a catalytic metal ion. Nature chemical biology, 13(5), 508–513. 10.1038/nchembio.2333

42. Bohdan, D. R., Voronina, V. V., Bujnicki, J. M., & Baulin, E. F. (2023). A comprehensive survey of long-range tertiary interactions and motifs in non-coding RNA structures. Nucleic acids research, 51(16), 8367–8382. 10.1093/nar/gkad605

43. Badepally, N. G., de Moura, T. R., Purta, E., Baulin, E. F., & Bujnicki, J. M. (2024). Cryo-EM structure of raiA ncRNA from Clostridium reveals a new RNA 3D fold. Journal of Molecular Biology, 436(23), 168833. 10.1016/j.jmb.2024.168833

44. Burley, S. K., Berman, H. M., Kleywegt, G. J., Markley, J. L., Nakamura, H., & Velankar, S. (2017). Protein Data Bank (PDB): the single global macromolecular structure archive. Protein crystallography: methods and protocols, 627–641. 10.1007/978-1-4939-7000-1_26

45. Kalvari, I., Nawrocki, E. P., Ontiveros-Palacios, N., Argasinska, J., Lamkiewicz, K., Marz, M., … & Petrov, A. I. (2021). Rfam 14: expanded coverage of metagenomic, viral and microRNA families. Nucleic acids research, 49(D1), D192–D200. 10.1093/nar/gkaa1047

46. Leontis, N. B., & Zirbel, C. L. (2012). Nonredundant 3D structure datasets for RNA knowledge extraction and benchmarking. RNA 3D structure analysis and prediction, 281–298. 10.1007/978-3-642-25740-7_13

47. Eustermann, S., Wu, W. F., Langelier, M. F., Yang, J. C., Easton, L. E., Riccio, A. A., … & Neuhaus, D. (2015). Structural basis of detection and signaling of DNA single-strand breaks by human PARP-1. Molecular cell, 60(5), 742–754. 10.1016/j.molcel.2015.10.032

48. Bonilla, S. L., Sherlock, M. E., MacFadden, A., & Kieft, J. S. (2021). A viral RNA hijacks host machinery using dynamic conformational changes of a tRNA-like structure. Science, 374(6570), 955–960. 10.1126/science.abe8526

49. Wang, B., Xu, W., & Yang, H. (2020). Structural basis of a Tn7-like transposase recruitment and DNA loading to CRISPR-Cas surveillance complex. Cell research, 30(2), 185–187. 10.1038/s41422-020-0274-0

50. Perederina, A., Li, D., Lee, H., Bator, C., Berezin, I., Hafenstein, S. L., & Krasilnikov, A. S. (2020). Cryo-EM structure of catalytic ribonucleoprotein complex RNase MRP. Nature communications, 11(1), 3474. 10.1038/s41467-020-17308-z

51. Pettersen, E. F., Goddard, T. D., Huang, C. C., Meng, E. C., Couch, G. S., Croll, T. I., … & Ferrin, T. E. (2021). UCSF ChimeraX: Structure visualization for researchers, educators, and developers. Protein science, 30(1), 70–82. 10.1002/pro.3943

52. Aleksandrova, E. V., Ma, C. X., Klepacki, D., Alizadeh, F., Vázquez-Laslop, N., Liang, J. H., … & Mankin, A. S. (2024). Macrolones target bacterial ribosomes and DNA gyrase and can evade resistance mechanisms. Nature chemical biology, 20(12), 1680–1690. 10.1038/s41589-024-01685-3

53. Petrov, A. I., Zirbel, C. L., & Leontis, N. B. (2013). Automated classification of RNA 3D motifs and the RNA 3D Motif Atlas. Rna, 19(10), 1327–1340. 10.1261/rna.039438.113

54. Lu, X. J., Bussemaker, H. J., & Olson, W. K. (2015). DSSR: an integrated software tool for dissecting the spatial structure of RNA. Nucleic acids research, 43(21), e142–e142. 10.1093/nar/gkv716

55. Leontis, N. B., & Westhof, E. (2001). Geometric nomenclature and classification of RNA base pairs. Rna, 7(4), 499–512. 10.1017/S1355838201002515

